# LysM receptors in *Coffea arabica*: identification, characterization, and gene expression in response to *Hemileia vastatrix*

**DOI:** 10.1101/2021.10.07.463563

**Authors:** Mariana de Lima Santos, Mário Lúcio Vilela de Resende, Bárbara Alves dos Santos Ciscon, Natália Chagas Freitas, Matheus Henrique de Brito Pereira, Tharyn Reichel, Sandra Marisa Mathioni

## Abstract

Pathogen-associated molecular patterns (PAMPs) are recognized by pattern recognition receptors (PRRs) localized on the host plant cell wall. These receptors activate a broad-spectrum and durable defense, which are desired characteristics for disease resistance in plant breeding programs. In this study, candidate sequences for PRRs with lysin motifs (LysM) were investigated in the *Coffea arabica* genome. For this, approaches based on the principle of sequence similarity, conservation of motifs and domains, phylogenetic analysis, and modulation of gene expression in response to *Hemileia vastatrix* were used. The candidate sequences for PRRs in *C. arabica* (*Ca1-LYP, Ca2-LYP, Ca1-CERK1, Ca2-CERK1, Ca-LYK4, Ca1-LYK5* and *Ca2-LYK5*) showed high similarity with the reference PRRs used: *Os-CEBiP, At-CERK1, At-LYK4* and *At-LYK5*. Moreover, the ectodomains of these sequences showed high identity or similarity with the reference sequences, indicating structural and functional conservation. The studied sequences are also phylogenetically related to the reference PRRs described in Arabidopsis, rice, and other plant species. All candidates for receptors had their expression induced after the inoculation with *H. vastatrix*, since the first time of sampling at 6 hours post-inoculation (hpi). At 24 hpi, there was a significant increase in expression, for most of the receptors evaluated, and at 48 hpi, a suppression. The results showed that the candidate sequences for PRRs in the *C. arabica* genome display high homology with fungal PRRs already described in the literature. Besides, they respond to pathogen inoculation and seem to be involved in the perception or signaling of fungal chitin, acting as receptors or coreceptors of this molecule. These findings represent an advance in the understanding of the basal immunity of this species.

## Introduction

The interaction between plants and pathogens can be understood as a co-evolutionary “molecular war,” in which each opponent uses their biological weapons as necessary, causing a successful infection by the pathogen or resistance in the host [1]. Currently, the study of pathogen perception by plants is divided into two lines. The first line is based on the recognition of conserved microbial molecules, denominated pathogen-associated molecular patterns (PAMPs), activating PAMP-triggered immunity (PTI). The second, on the other hand, recognizes the pathogen effectors by resistance proteins (R proteins), leading to effector-triggered immunity (ETI) [2,3].

The PAMPs recognition is performed by pattern recognition receptors (PRRs). These receptors are membrane proteins that usually have an extracellular domain involved in the perception of the ligand, the transmembrane or glycosylphosphatidylinositol (GPI) anchor domain that anchors the protein in the plasma membrane, and an intracellular kinase domain that is involved in the defense response signaling [4]. Adapted pathogens can suppress this first line of defense by secreting specific effectors. In response to this suppression, R proteins, encoded by resistance genes, recognize these effectors triggering ETI [5]. In spite of identifying different ligands, ETI and PTI lead to similar signaling pathways [6]. This signaling involves changes in calcium levels in the cytoplasm, production of reactive oxygen species (ROS) and signaling cascades involving protein kinases, MAPKs (mitogen-activated protein kinases) and CDPKs (calcium-dependent protein kinases) [7–10].

Comparing these two lines of defense, many studies indicate that the responses from the ETI occur more quickly and are more efficient than those from the PTI [11,12] since the former is associated with a hypersensitive response (HR), which involves programmed cell death and also systemic acquired response (SAR). For these reasons, the resistance conditioned by one or a few resistance genes has been the focus of breeding programs for several cultivated species. Nonetheless, the PTI is effective against pathogens, insects and parasitic plants and constitutes an important factor in non-host resistance [13,14]. In addition, it leads to a durable and broad-spectrum resistance [15,16]. The ETI, on the other hand, being characterized as a resistance against specific pathogens is quickly overcome, due to the emergence of new races of the pathogen [17].

Due to these defense characteristics, broad spectrum and durable, in which the PRRs are involved, currently they have been the target of studies aiming at a greater use of these receptors in plant breeding [16,18]. These studies focus on the possibility of combining (pyramiding) PRRs and increasing resistance to a broad spectrum of pathogens. The best characterized PRRs are the leucine-rich repeat receptor kinases (LRR-RKs). These receptors are involved in the recognition of bacterial structures. An example of this is *FLS2* (Flagellin sensing 2), which detects a conserved epitope of 22 amino acids, flg22, existing in the N-terminal region of the flagellin protein [18,19] and *EFR* (EF-Tu receptor), which detects the elf18 epitope, corresponding to the 18 conserved residues in the N-terminal region of the elongation factor Tu (EF-Tu) [20]. For fungi, well-described receptors are those that recognize chitin and have in common extracellular domains with lysin residues (Lys) [4,21], such as *CERK1* (chitin elicitor receptor kinase 1) [22], *CEBiP* (chitin elicitor binding protein) [23], *LYK4, LYK5* (LysM-containing receptor-like kinase 4 and 5) [24,25], *LYP4* and *LYP6* (LysM domain-containing protein 4 and 6) [26].

Genetic alterations in the PRRs that recognize both fungal and bacterial PAMPs reduce the plant ability to properly perceive and defend against pathogens. Gene knockouts such as *Os-CERK1* [21,22] and mutations in *At-LYK5* [24] lead to a loss of ability to respond to chitin and initiate defense responses to adapted pathogens. In addition, it allows some degree of disease progression by non-adapted pathogens, displaying failures in non-host resistance [15]. These studies demonstrate that the PTI and ETI form a continuum, which is necessary for a durable and efficient defense response [11]. Therefore, programs that seek to enable resistance to phytopathogens, with a focus on increasing the capacity of the recognition system, are successful by adding the PTI and ETI as the main strategy for obtaining resistant cultivars [15,27].

Few non-model plants, such as barley [28], apple [29,30] and mulberry [31], had PRRs characterized. *Coffea arabica* is an important coffee species cultivated in countries such as Brazil, Vietnam, Colombia, and Indonesia [32]. PAMP receptors have been scarcely studied in *Coffea spp*., therefore, it is crucial to identify the receptors that are present in their genome, and whether there is a response induced by the inoculation of pathogens, thus allowing the use of PRRs in coffee breeding programs.

The rust is the main coffee disease, causing severe losses in productivity in all regions where coffee is cultivated [33,34]. In Brazil, the biotrophic fungus *Hemileia vastatrix* Berk. & Br, the etiological agent of coffee rust, has caused damage since the 1970s [35,36]. In regions with favorable conditions for the pathogen, the decline in productivity can reach 50% [36]. To circumvent such damage, chemical control has been used, however, the use of tolerant or resistant cultivars is a viable alternative to reduce costs and possible environmental damage [33,37,38]. Therefore, the goals of this study were (i) to identify the pattern recognition receptors (PRRs) for fungi in the *C. arabica* genome, (ii) to characterize these sequences for protein domains and motifs and (iii) to analyze the gene expression of these PRRs in cultivars of *C. arabica* contrasting to rust resistance inoculated with *H. vastatrix*. The data obtained suggested that *C. arabica* has LysM receptors that act as fungal PAMP receptors, and that the expression of these receptors is stimulated after *H. vastatrix* inoculation. Our results contribute to the understanding and future employment of PRRs in coffee breeding programs.

## Materials and Methods

### Identification and characterization of specific PRRs for fungi in the *C. arabica* genome

The reference PRRs described in the literature for fungal PAMPs recognition in *Arabidopsis thaliana* and in *Oryza sativa* were selected: *At-CERK1, At-LYK4, At-LYK5* and *Os-CEBiP* (Table 1). To identify these receptors, the *C. arabica* genome (accession UCG-17, variety Geisha) sequenced by the University of California (UC Davis Coffee Genome Project) and partially available in the Phytozome database (https://phytozome.jgi.doe.gov/pz/portal.html) was used. The search was based on sequence similarity and domain conservation. For this, a BLASTp (Align Sequences Protein BLAST) with default parameters was performed in Phytozome. The *C. arabica* sequences returned by BLASTp were selected based on the following criteria: e-value ≤ 10^−5^, extracellular domain corresponding to the reference sequence used (Lysin motifs-LysM), and transmembrane or GPI anchor domain. The domains were analyzed using the SMART (http://smart.embl-heidelberg.de/), the TMHMM2.0 (http://www.cbs.dtu.dk/services/TMHMM/) and the PredGPI (http://gpcr.biocomp.unibo.it/predgpi/pred.htm).

**Table 1.**
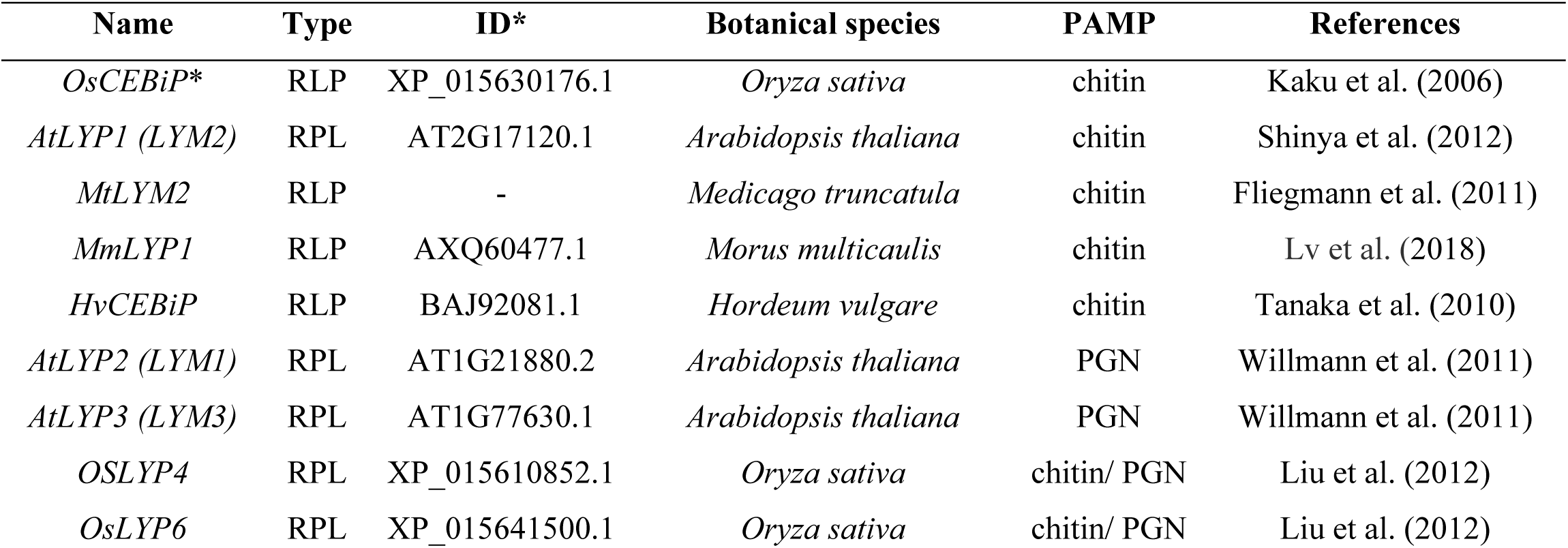
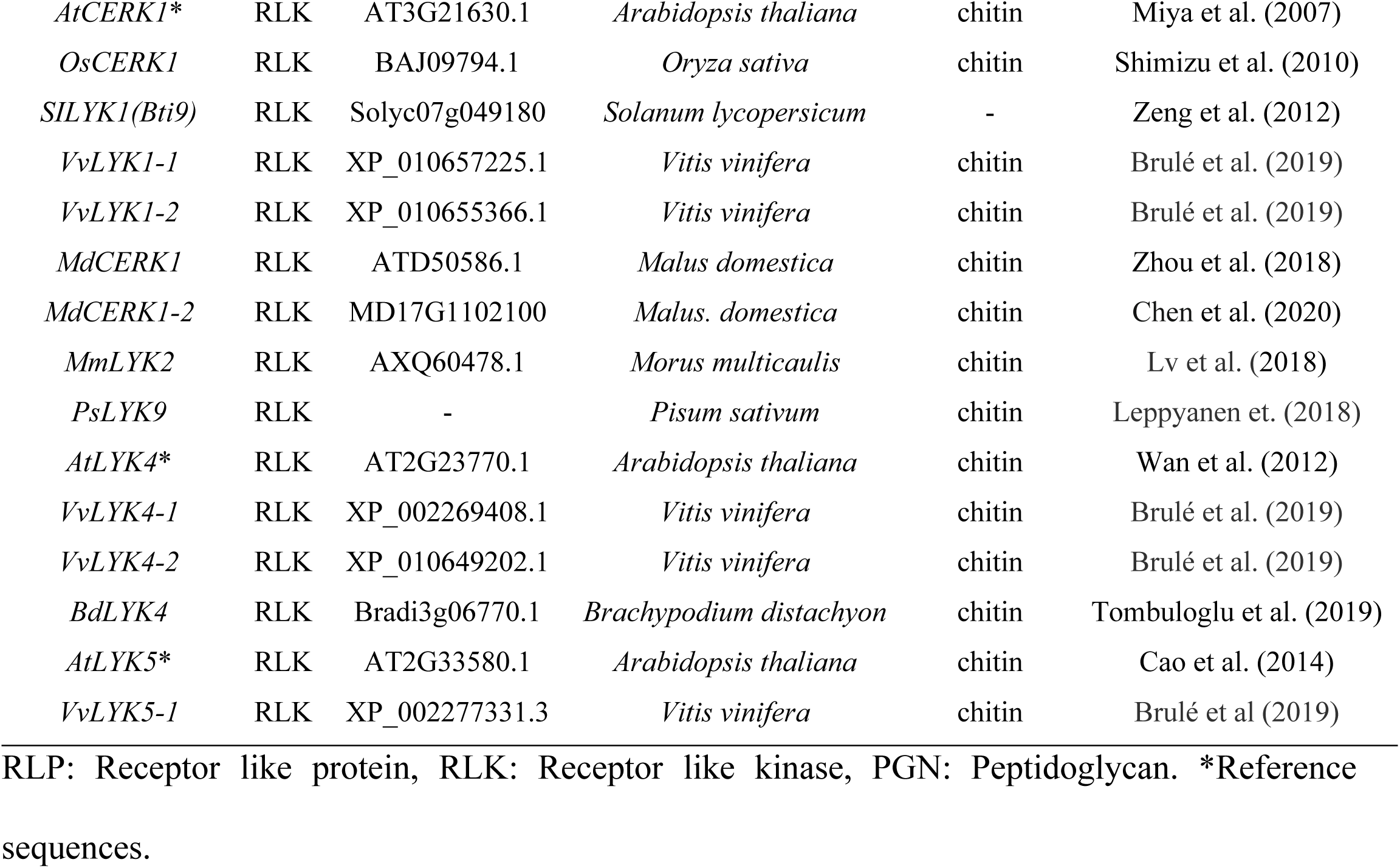
Reference PRRs and homologues.

| Name | Type | ID* | Botanical species | PAMP | References |
| --- | --- | --- | --- | --- | --- |
| *OsCEBiP\** | RLP | XP_015630176.1 | *Oryza sativa* | chitin | Kaku et al. (2006) |
| *AtLYP1 (LYM2)* | RPL | AT2G17120.1 | *Arabidopsis thaliana* | chitin | Shinya et al. (2012) |
| *MtLYM2* | RLP | - | *Medicago truncatula* | chitin | Fliegmann et al. (2011) |
| *MmLYP1* | RLP | AXQ60477.1 | *Morus multicaulis* | chitin | Lv et al. (2018) |
| *HvCEBiP* | RLP | BAJ92081.1 | *Hordeum vulgare* | chitin | Tanaka et al. (2010) |
| *AtLYP2 (LYM1)* | RPL | AT1G21880.2 | *Arabidopsis thaliana* | PGN | Willmann et al. (2011) |
| *AtLYP3 (LYM3)* | RPL | AT1G77630.1 | *Arabidopsis thaliana* | PGN | Willmann et al. (2011) |
| *OSLYP4* | RPL | XP_015610852.1 | *Oryza sativa* | chitin/ PGN | Liu et al. (2012) |
| *OsLYP6* | RPL | XP_015641500.1 | *Oryza sativa* | chitin/ PGN | Liu et al. (2012) |
| *AtCERK1\** | RLK | AT3G21630.1 | *Arabidopsis thaliana* | chitin | Miya et al. (2007) |
| *OsCERK1* | RLK | BAJ09794.1 | *Oryza sativa* | chitin | Shimizu et al. (2010) |
| *SILYK1(Bti9)* | RLK | Solyc07g049180 | *Solanum lycopersicum* | - | Zeng et al. (2012) |
| *VvLYK1-1* | RLK | XP_010657225.1 | *Vitis vinifera* | chitin | Brulé et al. (2019) |
| *VvLYK1-2* | RLK | XP_010655366.1 | *Vitis vinifera* | chitin | Brulé et al. (2019) |
| *MdCERK1* | RLK | ATD50586.1 | *Malus domestica* | chitin | Zhou et al. (2018) |
| *MdCERK1-2* | RLK | MD17G1102100 | *Malus. domestica* | chitin | Chen et al. (2020) |
| *MmLYK2* | RLK | AXQ60478.1 | *Morus multicaulis* | chitin | Lv et al. (2018) |
| *PsLYK9* | RLK | - | *Pisum sativum* | chitin | Leppyanen et. (2018) |
| *AtLYK4\** | RLK | AT2G23770.1 | *Arabidopsis thaliana* | chitin | Wan et al. (2012) |
| *VvLYK4-1* | RLK | XP_002269408.1 | *Vitis vinifera* | chitin | Brulé et al. (2019) |
| *VvLYK4-2* | RLK | XP_010649202.1 | *Vitis vinifera* | chitin | Brulé et al. (2019) |
| *BdLYK4* | RLK | Bradi3g06770.1 | *Brachypodium distachyon* | chitin | Tombuloglu et al. (2019) |
| *AtLYK5\** | RLK | AT2G33580.1 | *Arabidopsis thaliana* | chitin | Cao et al. (2014) |
| *VvLYK5-1* | RLK | XP_002277331.3 | *Vitis vinifera* | chitin | Brulé et al (2019) |

RLP: Receptor like protein, RLK: Receptor like kinase, PGN: Peptidoglycan. \*Reference sequences.

After selecting the sequences of *C. arabica*, they were again compared to the reference sequences by phylogenetic analysis. This analysis enabled to identify which peptide sequences had the greatest phylogenetic similarity to the reference PRRs, thus allowing the selection of candidate sequences. Additionally, considering that these PRRs present protein domains very close, a joint phylogenetic tree, with the candidate sequences in *C. arabica*, the reference PRRs and homologs (Table 1), was also created to confirm the separation of these groups and the homology of these sequences. The databases used to retrieve the reference sequences were: the GenBank from the National Center for Biotechnology Information (NCBI) sequence database, the Arabidopsis Information Resource (TAIR), the Sol Genomics Network, the Apple Genome and Epigenome, and Phytozome. The complete amino acid sequences were aligned by the CLC Genomics Workbench software version 11.0.1 (QIAGEN) (default parameters with very accurate) and the phylogenetic tree was generated by the Mega software version 10.1.8 [39] using the Maximum Likelihood method with a bootstrap of 1000 replications.

To characterize the extracellular regions of the candidate sequences, the lysin motifs (LysM) were used for multiple alignments between the candidate and reference sequences. The LysM motifs of each sequence were predicted by SMART using the extracellular region and aligned by the MAFFT program online version (https://mafft.cbrc.jp/alignment/server/) [40]. After the alignment, the visualization and calculation of the identity and similarity of each of the candidate sequences against the reference sequences were obtained by BioEdit version 7.2.5 [41].

Considering the fact that *C. arabica* is an allotetraploid (2n = 4x = 44 chromosomes), the result of the cross between *C. canephora* and *C. eugenioides* [42,43], the sequences selected as PRR candidates for the arabica coffee were also analyzed by BLASTp in the database of the NCBI (https://www.ncbi.nlm.nih.gov/) against each ancestral subgenome. This analysis aimed to verify the possible genomic origin of the studied PRRs.

### Primer design

The *C. arabica* sequences selected as candidates by the phylogenetic analysis were used for primer design. The primers were designed using the Primer Quest software and their quality was analyzed using the Oligo Analyzer software, both available online by IDT (Integrated DNA Technologies, USA). After the primers were designed, they were blasted (BLASTn - Standard Nucleotide BLAST) against the NCBI and Phytozome database (https://blast.ncbi.nlm.nih.gov/Blast.cgi) to attest their specificity through the identification of non-complementarity with nonspecific sequences.

### Fungal inoculum preparation

The inoculum used was obtained from leaves of *C. arabica* naturally infected with *H. vastatrix*. The pustules of these leaves were scraped and placed in microtubes, were frozen in liquid nitrogen, and stored in a freezer at −80ºC. To prepare the inoculum, the stored spores were submitted a 40ºC thermal shock for 10 min and were tested for viability by the spore germination test. The viability was verified by observing the spore germination in glass cavity slides. After preparing the suspension (1 × 10^6^ urediniospores/ml) for plant inoculation, three drops were transferred to glass cavity slides, which were incubated at 25°C for 48 hours. After the incubation, the spores were visualized under an optical microscope, so their germination could be observed (S1 Fig).

### Plant materials, experimental design, and inoculation

Aiming to analyze the gene expression of the PRR selected candidates, seedlings of four cultivars of *C. arabica* were used, being two rust susceptible cultivars, Catuaí Vermelho IAC 144 (CV) and Mundo Novo IAC 367-4 (MN), and two rust resistant, Aranãs RV (AR) and Iapar-59 (IP). The experiment was conducted in a randomized complete block design (RCBD) with three replicates and an experimental plot consisting of three plants. The treatments were arranged in a 2 × 3 × 4 factorial scheme, the factors being: condition (inoculated and not inoculated); evaluation times (06, 24 and 48 hours post-inoculation - hpi) and cultivars (Catuaí Vermelho IAC 144, Mundo Novo, Aranãs RV, and Iapar-59). The experiment was repeated twice independently.

Young plants (3-4 pairs of leaves) were inoculated in a growth chamber with a controlled environment (temperature of 22 ± 2°C, relative humidity of 90%) favoring the disease development. The suspension was sprayed on abaxial leaf surfaces and the inoculated plants were kept in the dark in a humid chamber according to a previously published methodology [44]. The control plants (sprayed with pure water only) were also sampled at all the evaluated time points. All the leaves collected were immediately frozen in liquid nitrogen and subsequently stored in a freezer at −80°C. After the treatment and sampling, the plants were kept in a greenhouse until the first symptoms and signs of the pathogen were seen to make sure the inoculation was effective (S2 Fig).

### RNA extraction and quantification

Following the RNA extraction, the samples were ground with liquid nitrogen until a fine powder was obtained. The ground material was stored in a ultrafreezer at −80°C until the RNA extraction was performed. The extraction was performed using the Plant RNA Purification Reagent (Thermo Fisher). Subsequently, the RNA was treated with DNase (RQ1 RNase-Free DNase, Promega) to remove any residual DNA in the sample. These procedures were performed according to manufacturer’s instructions. The integrity of the RNA was verified on 1% agarose gel and quantified on the NanoDrop One spectrophotometer (Thermo Fisher). All samples used showed a ratio reading 1.8-2.0 of absorbance at 260/280 nm and 260/230 nm for high-quality RNA.

### cDNA synthesis and RT-qPCR

An aliquot containing 1 μg of total RNA (treated with DNase) was used for cDNA synthesis using the High-Capacity cDNA Reverse Transcription Kit with RNase Inhibitor (Thermo Fisher). After the synthesis, the cDNA was diluted 5x and stored at −20 °C. The RT-qPCR were performed in the QuantStudio® 3 Real-Time PCR System (Applied Biosystems) using the SYBR® Green detection system. The amplification conditions were: 50°C for 2 min and 95°C for 10 min, 40 cycles: 95°C for 15 s, 60°C for 1 min and a final step of 95°C for 15 s (melting curve). The final reaction volume was 10μL contained the following components: 1.0 μL of cDNA (∼ 10 ng), 0.4 μL of each primer (forward and reverse) at a concentration of 10 μM (400 nM in the reaction), except for the *Ca2-CERK1* (Scaffold 2193.164 and 476.38), which used 0.2 μL (200 nM in the reaction), 5.0 μL of Platinum SYBR Green qPCR SuperMix-UDG with ROX (Thermo Fisher), and 3.4 μL of ultrapure water (free of nucleases).

For each of the three biological samples, technical triplicates were used and for each plate an inter-assay sample was used to ensure the reproducibility of the technique. The relative quantification was calculated according to the formula by Pfaffl, 2001 [45]. Referring to the data normalization, the expression stability of four reference genes was analyzed: protein 14-3-3 (*14-3-3*), glyceraldehyde-3-phosphate dehydrogenase (*GAPDH*), ribosomal protein 24S (*24S*) and factor elongation 1α (*EF1-α*) [46–49]. The efficiency correction of these genes in Cq values was performed by the GenEx Enterprise program (version 7.0) and the stability was verified by the RefFinder tool [50]. The two most stable genes were *14-3-3* and *GAPDH* (S3 Fig), which were used to normalize the transcription levels of the target genes. The samples with the lowest expression were used as calibrators. The MN 48 hpi was used as calibrator sample, except for the *Ca1-CERK1* (experiment 2), which was used the IP 48 hpi sample. The PCR amplification efficiencies and linear regression coefficients were determined using the LinRegPCR software version 2018.0 (Table 2) [51]. The average expression was obtained by the ratio of the sample inoculated with *H. vastatrix* compared to the average of the control treatment (without inoculation).

**Table 2.**
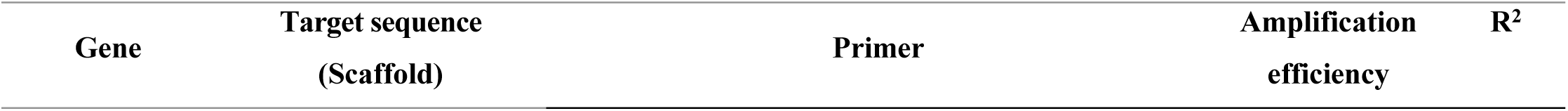
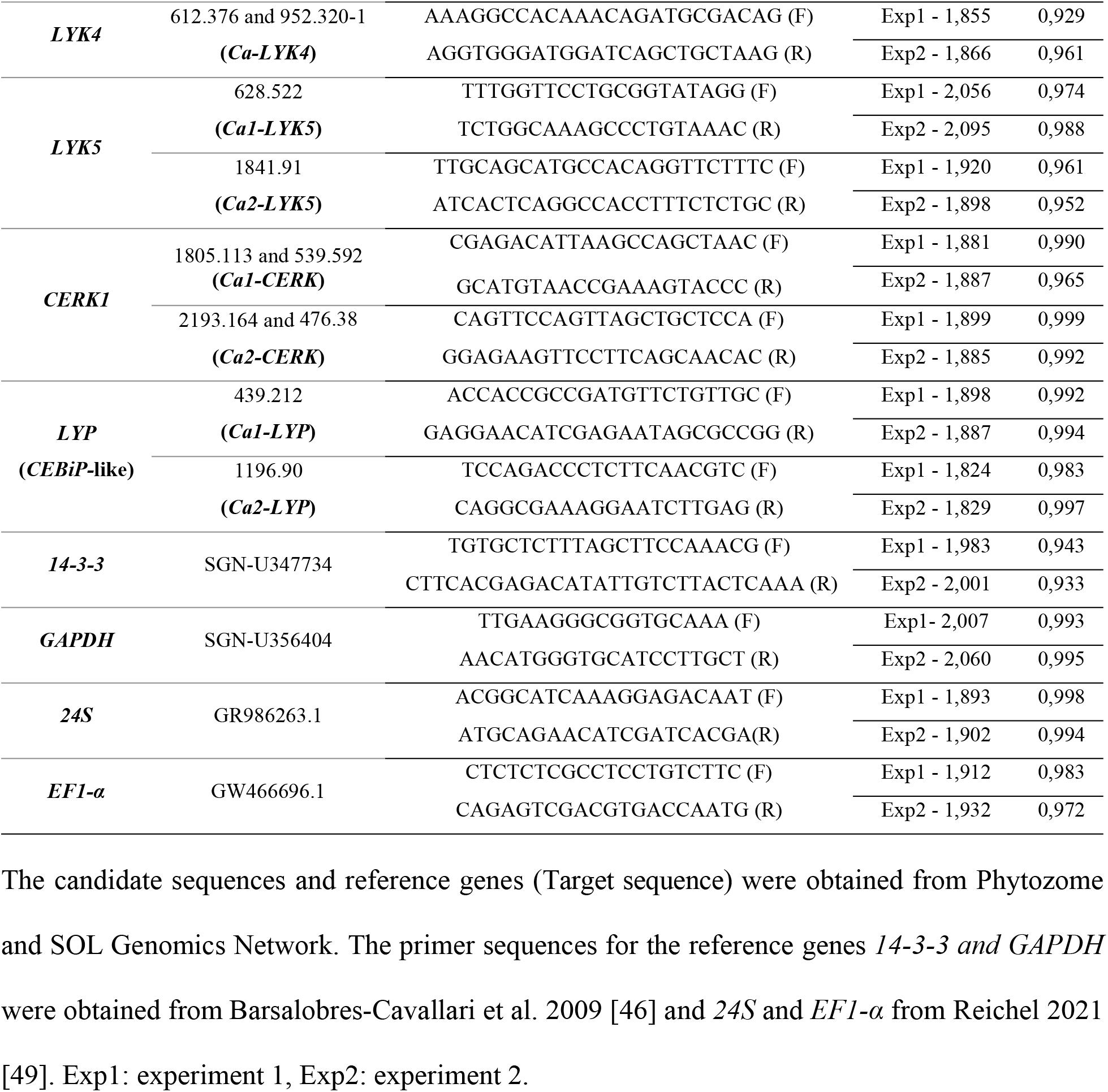
Sequence of primers used for candidate sequences of *C. arabica* PRRs and reference genes.

### Statistical analysis

The relative expression data of the two experiments were subjected to analysis of variance, using the following model:

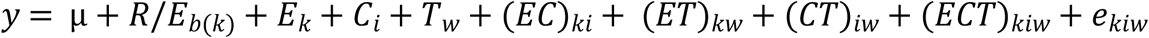

in which *R*/*E*_*b*(*k*)_ is the effect of block b within experiment k; *E*_*k*_ is the effect of experiment k, *C*_*i*_ is the effect of cultivar *i, T_w_* is the effect of time w, (*EC*)_*ki*_ is the effect of the interaction between experiment k and cultivar *i*, (*ET*)_*kw*_ is the effect of the interaction between experiment *k* and time w; (*CT*)_*iw*_ is the effect of the interaction between cultivar *i* and time w; (*ECT*)_*kiw*_ it is the effect of the interaction between experiment k cultivar *i* and time w; *e*_*kiw*_ is the effect of the experimental error, ∩ N(0, σ_e_^2^). Checks for outliers and of the assumptions of residuals from models were accomplished using diagnostic plots within the R software [52].

The interaction between cultivar and time was decomposed and the means between the levels of the factors were analyzed by Tukey’s test at 5% of probability. Data analysis was performed using the R software [52].

## Results

### Identification and characterization of specific fungal PRR in the *C. arabica* genome

The BLASTp analysis in Phytozome with the reference PRRs resulted in 4, 10, 12 and 14 sequences in the *C. arabica* genome for *Os-CEBiP, At-LYK5, At-CERK1* and *At-LYK4*, respectively (Fig 1 and S1 Table). These sequences were selected because they have e-value ≤ 10^−5,^ extracellular region containing lysin motif (LysM) and transmembrane domain or GPI-anchor. After the phylogenetic analysis, two candidate sequences were selected for *LYK4* (Scaffold 612.376 and 952.320) and *LYK5* (Scaffold 628.522 and 1841.91) (Fig 1B and 1D and S1Table) and four ones for *CERK1* (Scaffold 539.592, 1805.113, 2193.164 and 476.38) (Fig 1A and S1 Table). As the phylogenetic analysis for candidate sequences to the *CEBiP* protein did not result in a significant bootstrap (Fig 1C), other proteins belonging to the LYP clade (*CEBiP-like*) described in Arabidopsis and rice were included in a new analysis: *At-LYP1* (*At-CEBiP / LYM2*), *At-LYP2* (*LYM1*), *At-LYP3* (*LYM3*), *Os-LYP4* and *Os-LYP6* (Table 1).

**Fig 1.**
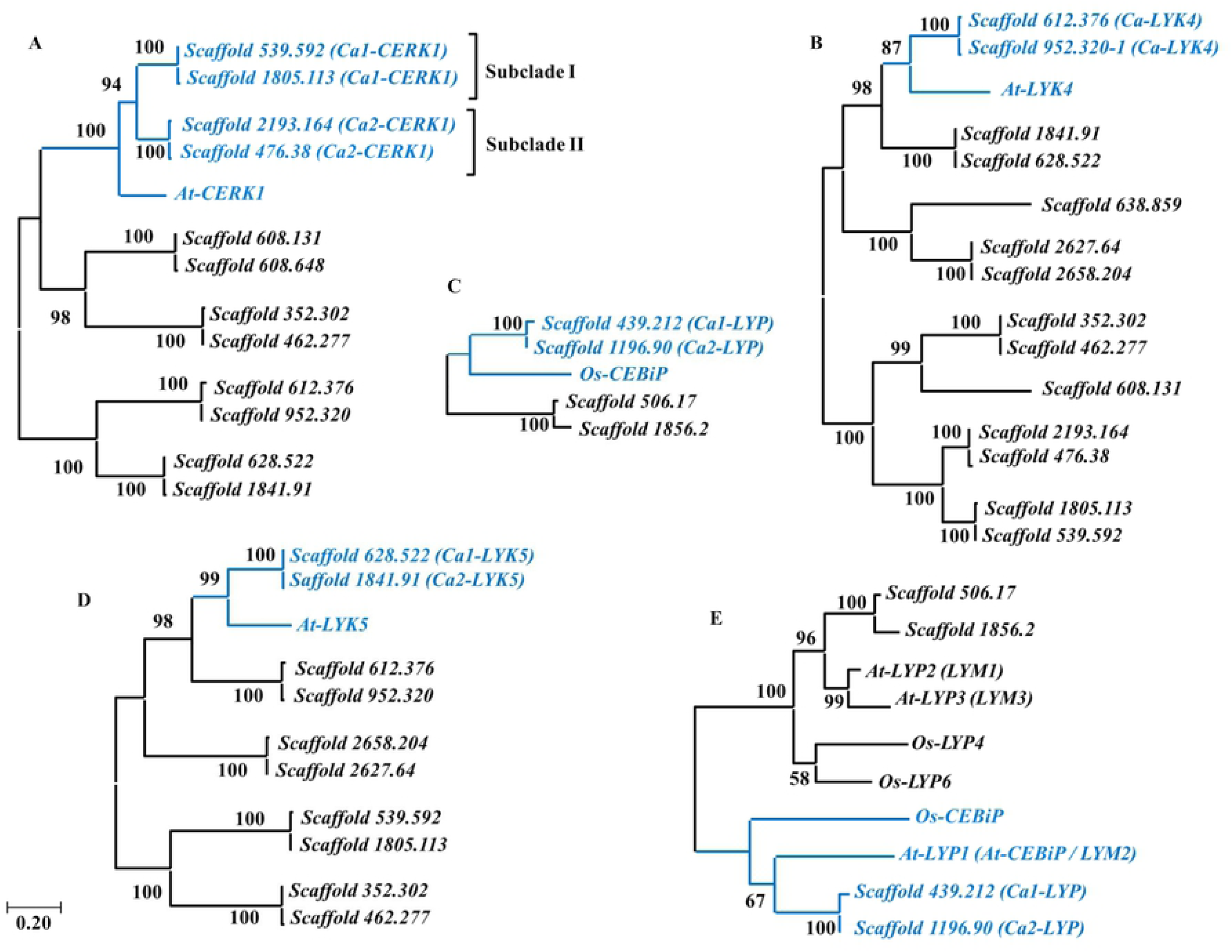
Phylogenetic analysis of the selected sequences for *C. arabica* by comparison with the reference PRRs. (A) *CERK1*, (B) *LYK4*, (C) *CEBiP*, (D) *LYK5*, (E) *CEBiP* and reference proteins belonging to the *LYP* (*CEBiP-*like) group. The phylogenetic trees were constructed with complete amino acid sequence alignments using the Maximum Likelihood method with a bootstrap of 1000 replications. The cluster clade of candidate sequences for *C. arabica* and reference sequences are highlighted in blue.

The new phylogenetic analysis for *CEBiP* (Fig 1E) showed two distinct clades. The clade one formed by the sequences Scaffold 506.17 and 1856.2, *At-LYP*2, *At-LYP3, Os-LYP4* and *Os-LYP6*, and the clade two formed by *Os-CEBiP, At-LYP1*, Scaffold 439.212 and 1196.90. As the Scaffold sequences 439.212 and 1196.90 showed greater similarity with the *Os-CEBiP* homologue in *A. thaliana* (*At-LYP1)*, they were selected as candidate sequences for the *CEBiP-like* (Fig 1 and S1 Table). Moreover, the *At-LYP2* (*LYM1*) and *At-LYP3* (*LYM3*), belonging to clade one, are described in the literature for their ability to recognize the peptidoglycan, a bacterial PAMP [53]. These sequences formed the nearest clade to the Scaffold 506.17 and 1856.2 sequences, substantiating the choice of the two *C. arabica* sequences belonging to clade two. The *Os-LYP4* and *Os-LYP6* that play a dual role, recognizing peptidoglycan and chitin [26], were not evaluated in this study.

All the domains found in the coffee candidate sequences correspond to the characteristic domains of the reference sequences. The description of these sequences such as identity and similarity in relation to the reference sequences as well as the gene size, the CDS and the number of exons, are shown in Table 3. The candidate sequences for *CERK1, LYK4* and *LYK5* have an extracellular LysM domain (with three LysM), a transmembrane domain, and an intracellular Ser/Thr kinase domain. The sequences selected as *CEBiP-like* have two lysin motifs and a predicted GPI-anchor. The characterization of these domains, motifs and protein sizes are shown in Fig 2.

**Table 3.** BLASTp and nucleotide characterization of candidate sequences in *C. arabica*.

| Candidate sequence | Identity (%) | Similarity (%) | Gene (pb) | Exons | CDS (pb) |
| --- | --- | --- | --- | --- | --- |
| *CERK1*-Scaffold_539.592 | 56.109 | 70.3 | 6082 | 13 | 2511 |
| *CERK1*-Scaffold_1805.113 | 55.145 | 69.0 | 4186 | 10 | 1815 |
| *CERK1*-Scaffold_2193.164 | 57.546 | 73.0 | 10180 | 12 | 1860 |
| *CERK1*-Scaffold_476.38 | 57.261 | 73.1 | 9921 | 12 | 1860 |
| *LYK4*-Scaffold_612.376 | 46.154 | 64.1 | 1935 | 1 | 1935 |
| *LYK4*-Scaffold_952.320 | 46.154 | 64.4 | 1935 | 1 | 1935 |
| *LYK5*-Scaffold_628.522 | 58.036 | 76.5 | 2031 | 1 | 2031 |
| *LYK5*-Scaffold_1841.91 | 58.631 | 76.5 | 2031 | 1 | 2031 |
| *LYP*-Scaffold_1196.90 | 42.258 | 56.1 | 2961 | 4 | 1098 |
| *LYP*-Scaffold_439.212 | 35.385 | 49.6 | 3598 | 5 | 954 |

Percentage of identity and similarity refer to BASTp analysis of candidate sequences against reference sequences *At-CERK1*, *At-LYK4*, *At-LYK5* e *LYP* (*CEBiP*- like). Candidate sequences were obtained from Phytozome database.

**Fig 2.**
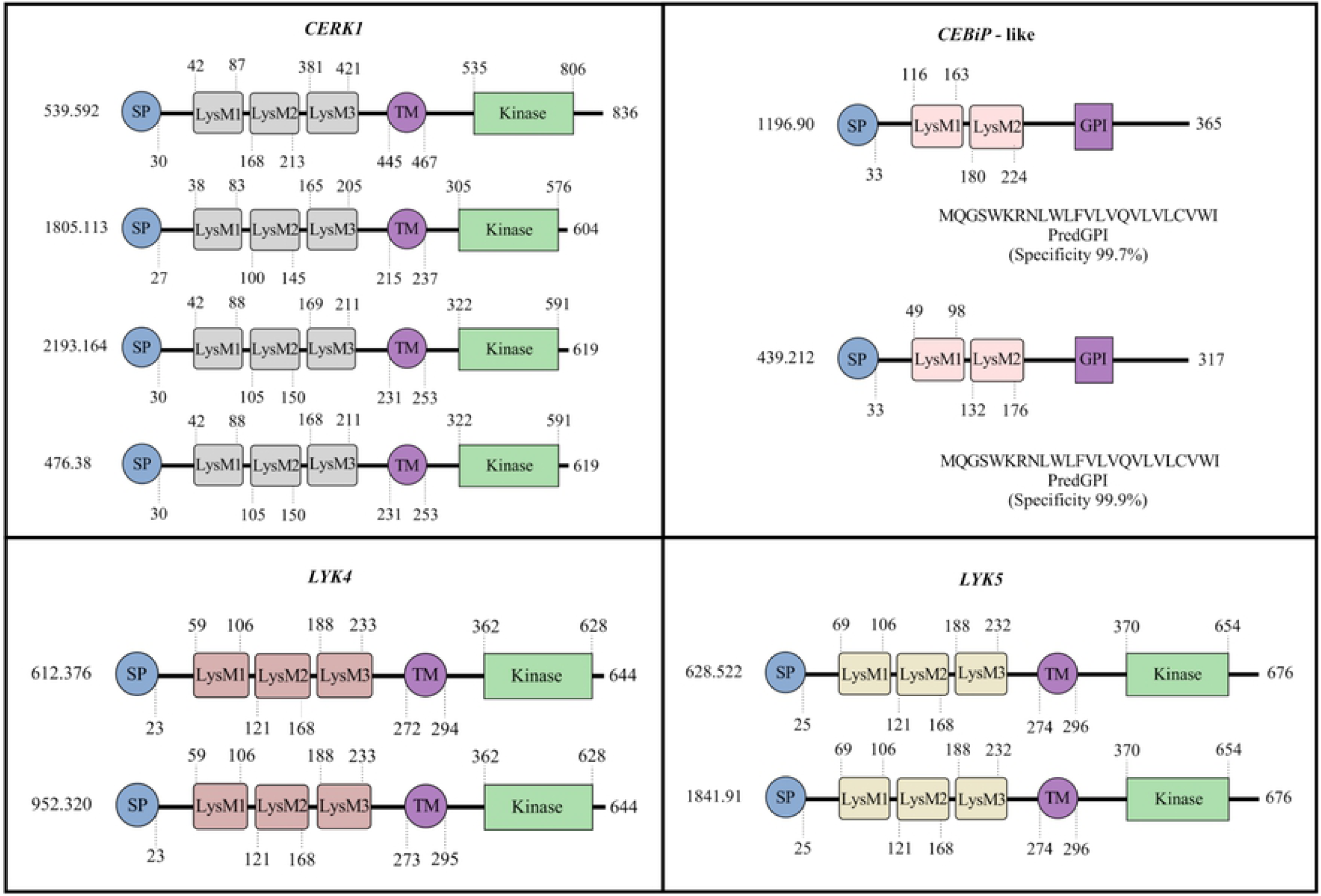
Protein characterization of the candidate sequences for *CERK1, LYK4, LYK5*, and *CEBiP*-like in *C. arabica*. The signal peptide positions, lysin motifs (LysM) and transmembrane domains were identified by SMART, and the GPI anchor by PredGPI. The domains positions are represented by numbers at the beginning and end of each domain. Concerning the *CEBiP*-like candidate sequences, the putative signal sequences for the GPI anchor and their specificities are shown. The numbers at the beginning of each sequence represents the scaffold (candidate sequence in *C. arabica*). The numbers at the end of each sequence represents the size of the proteins in number of amino acids. SP: signal peptide, LysM: lysin motifs identified as 1,2 e 3, TM: transmembrane domain, GPI: GPI-anchor.

The extracellular lysin motif regions (LysM1, LysM2 and LysM3) for these sequences ranged from 38 to 49 aa. The multiple alignments of these regions with the reference proteins showed high residue conservation but varied among the studied receptors (Fig 3). Out of eleven residues described as important for the chitin oligomer binding function in *At-CERK1* [54,55], eight ones displayed identity or similarity with the candidate sequences in *C. arabica*. For *Os-CEBiP*, from nine described [56], only three were present. In *At-LYK5*, only one of three described [24] showed similarity with *C. arabica* sequences. The tyrosine (Tyr) residue, located at position 128 in *At-LYK5*, considered as the fourth chitin-binding residue for this receptor, was not analyzed, as it is present between the LysM1 and LysM2 motifs, a region that was not analyzed in the alignment.

**Fig 3.**
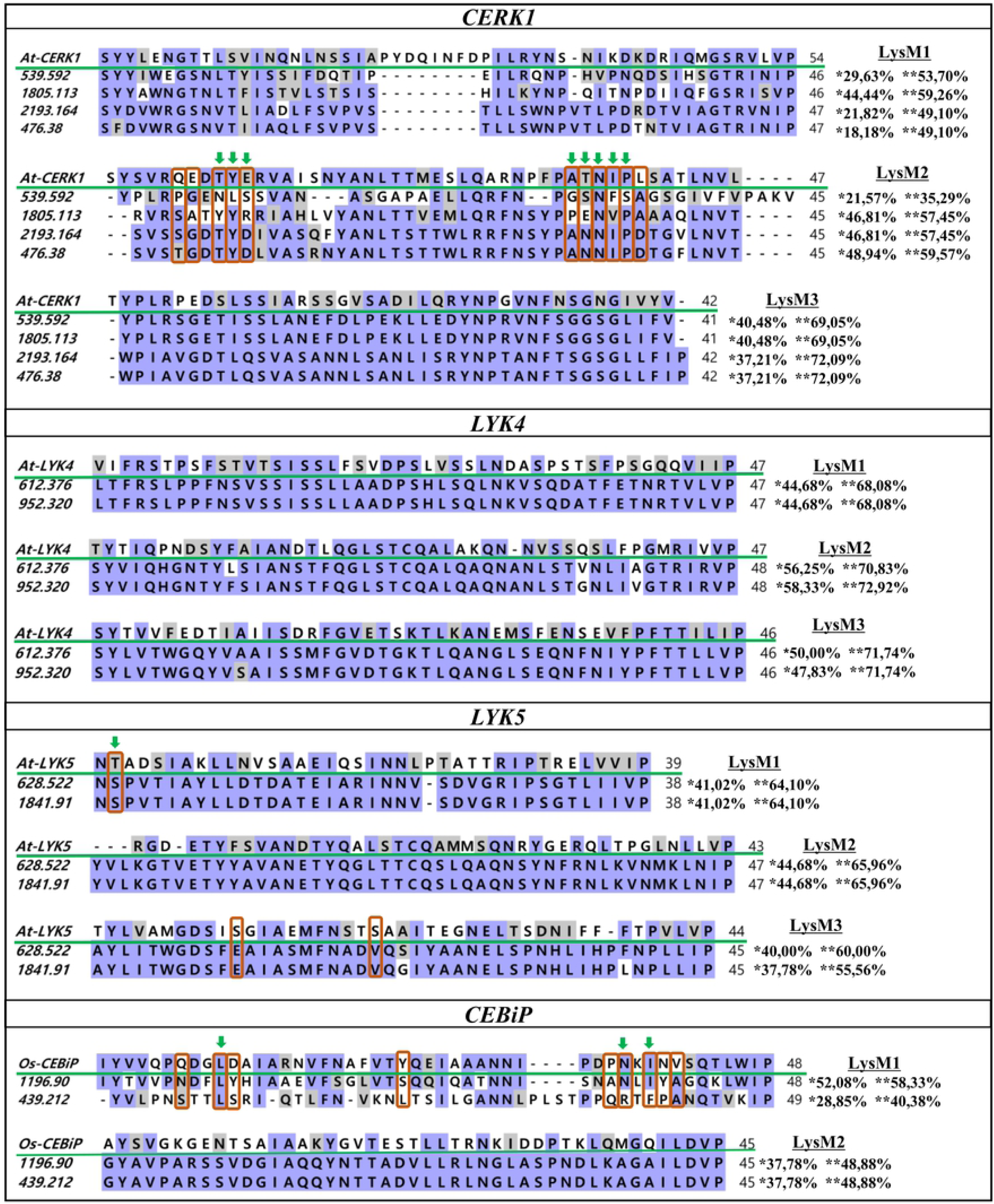
Alignment of the LysM motifs between reference sequences and candidate sequences in *C. arabica*. The LysM motif sequences were aligned using MAFFT and visualized by BioEdit. The numbers at the beginning of each sequence represents the scaffold (candidate sequence in *C. arabica*). The green line highlights the reference sequence. The purple and gray shading represent identical and similar amino acids, respectively. The percentages of identity and similarity between candidate sequences and references are indicated by * and **, respectively. In red are the critical residues that bind to chitin and the green arrows indicate residues identical or similar to these regions present in the candidate sequences in *C. arabica*. The numbers at the end of each sequence represent the size of the LysM motifs in number of amino acids.

### Joint phylogenetic analysis and BLASTp against the genomes of *C. canephora* and *C. eugenioides*

A joint phylogenetic tree was created to verify whether the candidate sequences would form distinct clades, including the reference sequences used. This tree was composed of the selected candidate sequences for PRRs in *C. arabica*, the reference sequences used to search for these PRRs in coffee (*At-CERK1, At-LYK4, At-LYK5* and *Os-CEBiP*) and homologs of these proteins described experimentally in the literature (Table 1). This analysis formed four clades that separated the candidate sequences in coffee with the respective reference proteins used, confirming their phylogenetic relationships (Fig 4).

**Fig 4.**
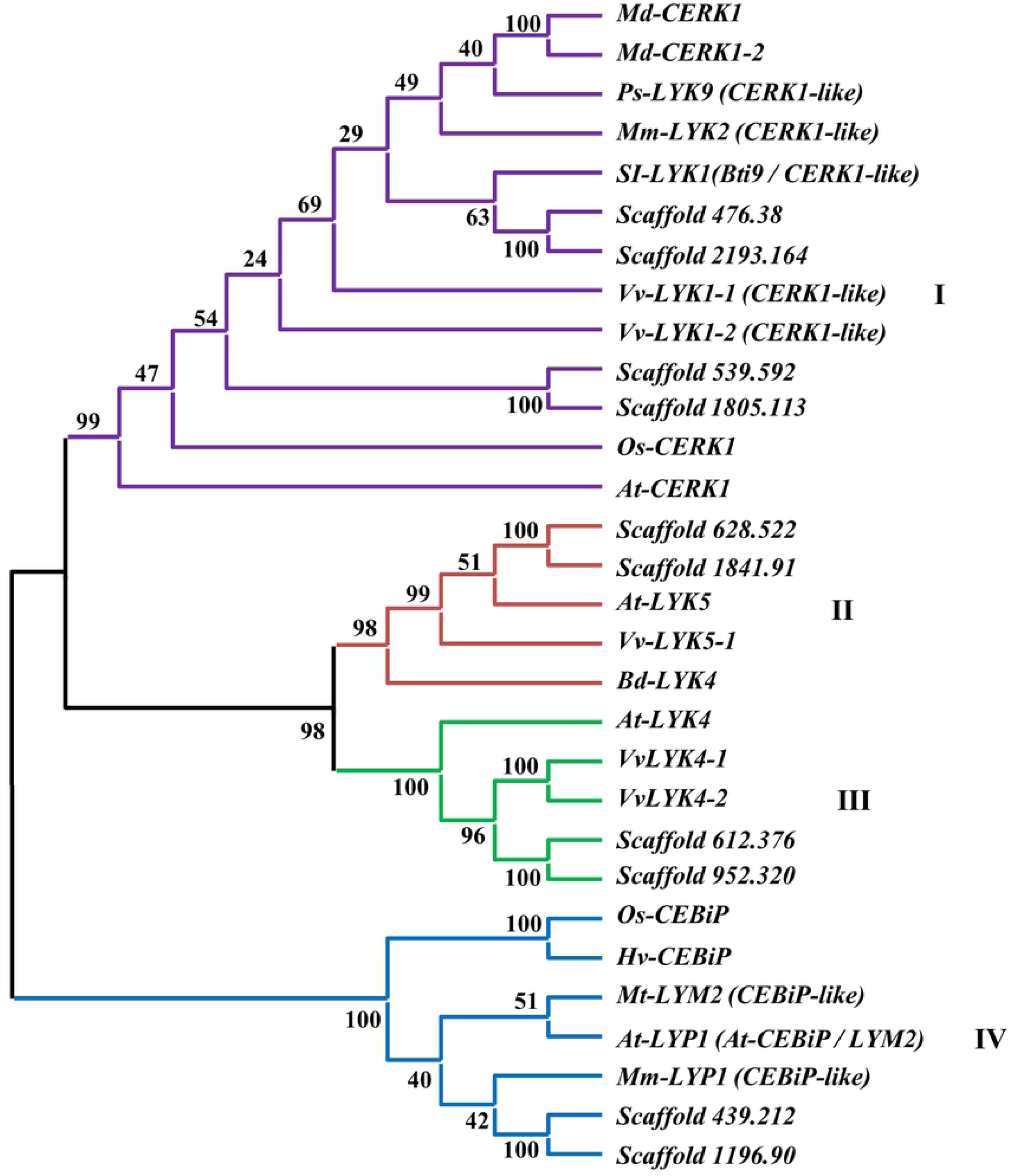
Joint phylogenetic analysis of candidate sequences in *C. arabica*, reference sequences and homologs described experimentally. The phylogenetic tree was constructed with alignments of complete amino acid sequences using the Maximum Likelihood method with a bootstrap of 1000 repetition. The *CERK1, LYK5, LYK*4 and *CEBiP*-Like clades are highlighted in different colors: I-purple, II-red, III-green and IV-blue.

The clade I was composed of Scaffold 539.592, 1805.113, 2193.164 and 476.38, *At-CERK1* and their homologs *Md-CERK1, Md-CERK1-2, Mm-LYK2* (*CERK1-*like), *Ps-LYK9, SI-LYK1* (*Bti9*), *Vv-LYK1-1, Vv-LYK1-2* and *Os-CERK1*. In this clade, the candidate sequences in coffee, Scaffolds 476.38 and 2193.164 are closest to the homologs of *At-CERK1* in tomato, *SI-LYK1* (*Bti9*), while the Scaffold 539.592 and 1805.113 sequences, formed a more distant subclade. Clades II and III belonging to *LYK4* and *LYK5* formed closer clades. The coffee sequences were grouped more closely to the *LYK4* homologues in grape and for the *LYK5* they formed a subclade with the reference sequence *At-LYK5* and its homolog also in grape (*Vv-LYK5-1*). In clade IV, belonging to the *CEBiP* cluster, it was observed that candidate sequences in coffee were significantly grouped with the *Os-CEBiP* homologs.

The BLASTp analysis in the NCBI database against the genomes of *C. canephora* and *C. eugenioides* showed that seven candidate sequences for PRRs in coffee have the highest percentage of identity with sequences belonging to the genome of *C. eugenioides* and only two (*LYK4*-Scaffold 612. 376 and *LYP*-Scaffold 1196.90) showing greater identity with the *C. canephora* sequences (Table 4).

**Table 4.**
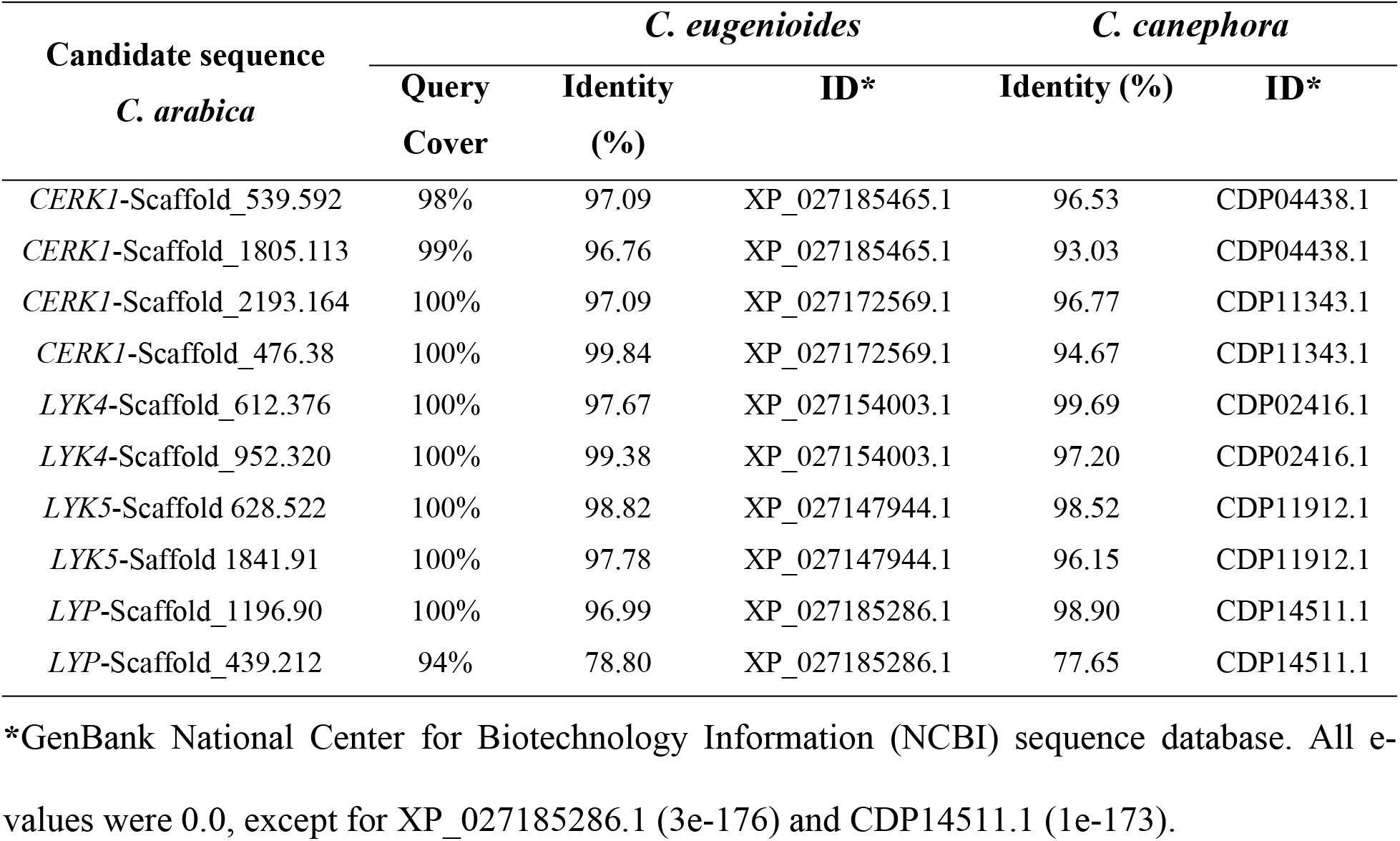
BLASTp analysis of candidate sequences in *C. arabica* against the genomes of *C. eugenioides* and *C. canephora*.

### Primer design

The four sequences selected as candidates for *CERK1* in the *C. arabica* genome by phylogenetic analysis formed two distinct subclades (Fig 1A). The subclade I formed by the Scaffold 539.592 and Scaffold 1805.113 sequences and the subclade II formed by the Scaffold 2193.164 and Scaffold 476.38 sequences. The coding sequences (CDS) of subclade I showed an 71.33% identity, with the 1805.113 sequence presenting a smaller CDS (1815bp) and shared almost entirely with the Scaffold 539.592 sequence. The Scaffold 539.592 sequence, on the other hand, presents a larger CDS (2511 bp) with two regions that are not present in 1805.113 (S4 Fig). The Scaffold 2193.164 and 476.38 showed CDS of the same size (1860bp) and an identity 98.28% (S5 Fig). For the primer design in the gene expression analysis, the formation of these two subclades was considered, thus using a pair of primers for each of the formed subclades. They were named *Ca1-CERK1* and *Ca2-CERK1* respectively and are referred to as such in the gene expression analysis (Table 2).

Concerning the *LYK4* candidate sequences (Scaffold 612.376 and 952.320), a primer pair was also designed for both candidate sequences. These showed a 98.45% identity (S6 Fig) and were named as *Ca-LYK4*. Regarding the candidate sequences *LYK5* (Scaffold 628.522 and Scaffold 1841.91) and *LYP* (*CEBiP-*Like) (Scaffold 439,212 and 1196.90), a primer pair was designed for each sequence separately and they are referred to as *Ca1-LYK5, Ca2-LYK5, Ca1-LYP, Ca2-LYP*, respectively (Table 2).

### Transcriptional response of candidate receptors in *C. arabica*

To verify the transcriptional responses of the candidate sequences to the PRRs in *C. arabica*, four cultivars with contrasting rust resistance levels were inoculated with *H. vastatrix*. The inoculum used displayed viability in both tests: the one with the glass cavity slides (S1 Fig) and the other about the ability to cause the disease symptoms and signs in susceptible cultivars CV and MN (S2 Fig). The resistant cultivars AR and IP, presented no symptoms or signs of the disease. The fungal inoculation induced the expression of all candidate receptors in all cultivars and studied time points. To a greater or lesser degree, there was an increase in expression from 6 hpi (Fig 5), with the peak varying between 6 and 24 hpi, followed by a decrease at 48 hpi.

**Fig 5.**
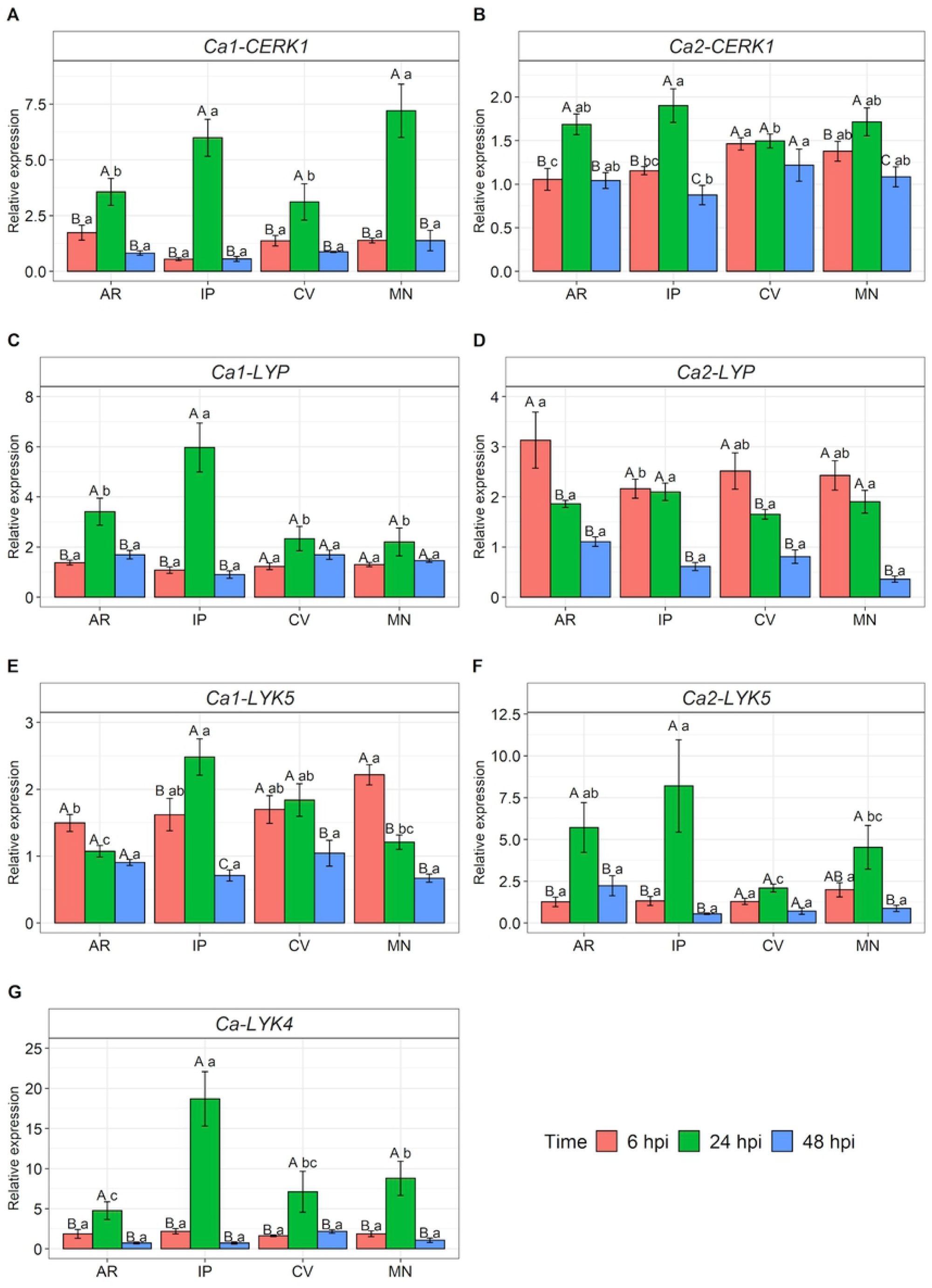
Relative expression of candidate genes for *CERK1, LYP* (CEBiP-like), *LYK5* and *LYK4* in *C. arabica*. (A) *Ca1-CERK1*, (B) *Ca2-CERK1*, (C) *Ca1-LYP*, (D) *Ca2-LYP*, (E) *Ca1-LYK5*, (F) *Ca2-LYK5*, (G) *Ca-LYK4*. Candidate genes were evaluated in *C. arabica* leaves at 6, 24 and 48 hours post-inoculation (hpi) with *H. vastatrix*. The average of relative expression was obtained by the ratio between the means of inoculated and control (not inoculated). Capital letters represent the statistical analysis of the times for each cultivar and lower letters between cultivars. Means followed by the same letter are not differentiated by Tukey’s test at 5% probability. The data shown represents experiments 1 and 2. MN: Mundo Novo, CV: Catuaí Vermelho IAC 144, AR: Aranãs RV, IP: IAPAR-59.

The two groups of candidate sequences for *CERK1* showed different expression profiles (Fig 5A and 5B) at 24 hpi. The *Ca1-CERK1* had higher expression than *Ca2-CERK1*. Concerning the former, the expression rate was seven times higher than that of the control in cultivar MN, regarding the latter, the highest value did not reach twice as much for IP. When the time expression levels were analyzed for each cultivar in the two groups (Fig 5A and 5B), there was a significant difference for 24 hpi, except for CV *Ca2-CERK1*. Regarding the *Ca1-CERK1*, the analysis between cultivars (Fig 5A) showed that IP and MN displayed approximately 6- and 7-fold higher expression levels at 24 hpi, respectively, demonstrating significant differences compared to AR and CV. Respecting 6 and 48 hpi, there were no significant differences. Concerning *Ca2-CERK1* (Fig 5B), the analysis between cultivars showed that at 6 hpi it was the most expressed in CV and MN. At 24 hpi, the highest expression was in IP, and at 48 hpi the same cultivar showed a reduction in its expression, which was the least expressed among the cultivars.

A similar profile to *CERK1* was observed for the sequences studied as candidates for *LYP* and *LYK5* (Fig 5 C - 5 F). The *Ca1-LYP* and *Ca2-LYK5* obtained cultivars with higher expression levels at 24 hpi than *Ca1-LYK5* and *Ca2-LYP*, however, for these genes, the candidate sequences were studied apart. Considering *Ca1-LYP* and *Ca2-LYP* (Fig 5C and 5D), the expression patterns were different at 6 and 24 hpi. The *Ca1-LYP* expression levels did not reach twice as much compared to the control at 6 hpi, while for *Ca2-LYP* the highest averages were observed at that time. Moreover, regarding the *Ca1-LYP*, all cultivars showed an expression above twofold higher at 24 hpi. Therefore, the greatest inductions for *Ca2-LYP* occurred at 6 hpi while for *Ca1-LYP* they happened later at 24 hpi.

The expression differences in time for each cultivar considering *Ca1-LYP* (Fig 5C) showed that AR and IP have significant differences at 24 hpi, which did not occur in CV and MN. The analysis between cultivars showed that at 6 hpi and 48 hpi there were no differences, but that at 24 hpi, IP was the cultivar that showed the highest expression, reaching 6-fold higher. Considering *Ca2-LYP* (Fig 5 D), AR and CV showed higher expressions at 6 hpi. For IP and MN, the largest expression occurred at 6 and 24 hpi, with no difference between these times. The analysis between cultivars showed that at 6 hpi, AR obtained the highest expression while IP presented the lowest expression. On the other hand, at 24 and 48 hpi, there were no differences between cultivars. However, it was found that 48 hpi was the time with the lowest average observed, within and between cultivars.

Regarding *Ca1-LYK5* (Fig 5E), there was a difference between the times for all cultivars, except for AR. The MN cultivar had the highest average at 6 hpi, while IP obtained the highest at 24 hpi. For the cultivar CV, there were no differences between these times, only at 48 hpi. Concerning the analysis between cultivars, the MN obtained the highest average at 6 hpi and IP at 24 hpi. At 48 hpi, there were no differences between cultivars and this time presented the lowest average for all. Referring to *Ca2-LYK5* (Fig 5F), all cultivars showed differences between the evaluated times, except for CV. The AR and IP cultivars showed significant differences in averages at 24 hpi compared to the ones at 6 and 48 hpi, coming to express about six and eight times more than the control, respectively. Regarding MN, the highest average was also detected at 24 hpi, but this did not differ statistically from 6 hpi, only from 48 hpi. For the times between cultivars, there were differences only in 24 hpi, with AR and IP having the highest expression.

The values for *Ca-LYK4* were the result of a single primer pair designed for two candidate sequences. In this receptor, the expression levels at 24 hpi differed within and between the cultivars evaluated. The IP cultivar obtained the highest average expression, reaching almost 19 times higher than that of the control, followed by MN, which expressed ninefold higher. The lowest averages for that time were observed for CV and AR, with an expression seven- and sixfold higher, respectively. For 6 and 48 hpi there was no difference within and between cultivars, the averages for those times reached at most twice as much.

## Discussion

### Fungal PRRs in the *C. arabica* genome

Understanding basal immunity has been the focus of several studies with the purpose of identifying the mechanisms governing this line of defense, enabling its use as another tool in the search for plant resistance to pathogens [18]. The description of the reference PRRs and studies of the modulation of their gene expression in response to *H. vastatrix*, one of the most devastating pathogens in coffee trees, presents an advance for understanding this crop basal immunity. In the present study, fungal PRR candidate sequences well described in the literature for model plants such as Arabidopsis and rice were studied in *C. arabica*. We observed that there is more than one candidate sequence for each receptor studied, which may be the result of the ploidy of this species or duplication of these receptors, a common mechanism in plant genomes [57]. Four candidate sequences for *CERK1* and two for *LYK5* in *C. arabica* presented higher percentages of identity compared to sequences of *C. eugenioides*, which may indicate duplications of this receptor in both species. Referring to *LYK4* and *LYP* (*CEBiP*-like), each candidate sequence showed greater identity with sequences of *C. eugenioides* or *C. canephora*. Therefore, it is possible to infer that those genes may have come from these genomes (Table 4).

The size of the CDS and the organization of exons demonstrated that the genes encoding *LYK4* and *LYK5* candidate proteins in *C. arabica* do not have introns, and the coding sequences are the result of a single exon. In fact, when compared to *CERK1* or *CEBiP*, these receptors are closer to each other in phylogenetic analysis. These results (Fig 4) corroborates with others described in the literature [54,58] and shows a greater evolutionary relationship between these receptors. Homologs of the *At-LYK4* and *At-LYK5* in many plant species have no introns and the coding region is the result of a single exon [25,58–61]. Regarding the LysM receptors homologous to *At-CERK1*, the CDS region mostly presents around 1800 bp with ten to twelve exons [29,54,62], which is likewise with the size of the CDS and number of exons found for the *CERK1* candidate sequences in coffee, except for the Scaffold 539.592, which presents a larger coding region, with 2511bp and 13 exons. However, this number of thirteen exons has also been found in *Ps-LYK9*, a *CERK1-like* gene in peas, which is involved in the control of plant immunity and symbiosis formation [62].

Regarding the genes LYPs (Receptor-like proteins or RLPs) such as *Os-CEBiP*, the number of exons reported is more variable from two to six [23,58,63]. In *C. arabica*, Scaffold 1196.90 and 439.212 presented four and five, respectively. The structural pattern of genes, such as the distribution of introns or exons in gene families, reinforces the ortholog identification between sequences since these are almost conserved among all orthologous. Minor differences may be due to evolutionary changes or errors in gene structure predictions [59].

### Characterization of domains and motifs (LysM)

Proteins classified as LYKs (Receptor-like kinases or RLKs) are composed of lysin motifs (LysM)-containing ectodomains, a transmembrane domain and an intracellular kinase. LYP proteins (RLPs), on the other hand, present LysM ectodomain, but without intracellular kinase and can be anchored to the plasma membrane by a transmembrane domain or GPI-anchor [58,64]. The *At-CERK1, At-LYK4* and *At-LYK5* contain three extracellular LysM motifs, a transmembrane domain and intracellular kinase, while *Os-CEBiP* has two extracellular LysM motifs and GPI anchor [22–24]. The SMART and PredGPI analysis predicted that the amino acid sequences of the PRRs studied in *C. arabica* present a signal peptide, extracellular LysM motifs, a transmembrane domain, or a putative signal sequence for the GPI anchor, besides the presence or absence of intracellular kinase. These characteristics differentiate them into LYKs (Ca1 and 2 *CERK1*, Ca1 and 2 *LYK5* and *Ca-LYK4*) and LYPs (Ca1 and 2 *LYP*) (Fig 2) and suggest that they all act as membrane receptors.

As a result of the organization of the domains, these proteins have different protein sizes. LYKs are generally larger than LYPs because they have an additional kinase domain. Protein sequences reported for these classes of receptors are around 500 or 600 and 300 or 400 aa respectively [23,58,65]. Candidate sequences in coffee have equivalent sizes, except for Scaffold 539.592 with 836aa, which may be a consequence of the size of the coding region.

The PRR extracellular region varies in plant with sizes from 35 to 50 aa [57,58]. These regions define the type of recognized PAMP and its binding affinity in addition to the interaction between receptors and coreceptors [66]. Differences in the chitin-binding properties between *At/Os-CERK1* ectodomains show variation in the performance of these receptors in Arabidopsis and rice. *At-CERK1*and *At-LYK5*, for instance, bind directly to chitin through their ectodomains containing LysM motifs with different affinities to the ligand, while *At-LYK4* appears to be a co-receptor [22,24,67]. In rice, *Os-CERK1* does not bind to chitooligosaccharides and the heterodimerization between *Os-CERK1* and *Os-CEBiP* is necessary for the innate immune response in this species [21,68]. Distinction in the role of these receptors suggests that plants use different chitin binding and signaling strategies [25,69].

In *C. arabica*, this region varied from 38 to 49 aa and the candidate sequences showed a high degree of identity and/or similarity with the reference LysM sequences used, indicating a conserved extracellular structure [54,56]. For *CERK1*, eight residues reported as important for chitin binding in Arabidopsis are present in the Scaffold 2193.164 and Scaffold 476.38 sequences (seven identical and one similar), suggesting that they can bind chitin. However, complementary data are still needed to clarify which would be the primary receptor and co-receptor of the innate immunity in this species, and further studies of chitin-receptor and receptor-receptor interaction are required.

### Joint phylogenetic analysis

PRRs are conserved in several plant species [59].This conservation indicates a fundamental importance of the PAMP recognition system [26]. The joint phylogenetic analysis showed that the sequences selected as candidates for *CERK1* in coffee, were highly related to *Md-CERK1, Md-CERK1-2, Ps-LYK9, Mm-LYK2, Vv-LYK1-1, Vv-LYK1-2, Os-CERK1* and *At-CERK* (Fig 4). All of these proteins have been described as being involved in the defense against fungal pathogens [21,22,29–31,54,62], suggesting that the studied sequences also participate in the defense responses against this group of phytopathogens. Among the species compared, tomato and grape have greater evolutionary proximity to coffee. *Bti9* (*Sl-LYK1*), a *CERK1* homolog in tomato, which grouped more closely to the Scaffold 2193.164 and 476.38 sequences (*Ca2-CERK1*) in this clade, presents an identity of 58.6% with *At-CERK* [70]. Candidate sequences in coffee, however, showed around 57% of identity (Table 3).

The *Bti9* (*Sl-LYK1*) in tomato interacts with *AvrPtoB*, effector in *Pseudomonas syringae*. The kinase region of this protein is the target and this results in blocking the PTI signaling [70]. Despite being described as a bacterial effector target, the study by Zeng et al., 2012 [70] or later reports by Xin and He, 2013 [71] did not describe the interaction of this protein with chitin or the transcriptional profiles regarding the response to fungal pathogens. Nonetheless, *Bti9* is a membrane receptor with extracellular LysM motifs and high homology to *At-CERK1*. Furthermore, the *At/Os-CERK1*, besides playing a role as a receptor for fungal PAMPs, also participates as a co-receptor for PRRs in bacterial recognition [53,72], which demonstrates the multiple functions of this receptor and turns it into a possible target of bacterial and fungal effectors that suppress PTI.

The Ca1 and 2 *LYK 4* and 5, clades II and III, were grouped to grape receptors *Vv-LYK4-1/2* and *Vv-LYK5-1* (Fig 4). These were shown to be highly expressed during infection by *Botrytis cinerea* in grapevine fruits [50]. The clustering of *Bd-LYK4* in this clade corroborates the results presented by Tombuloglu et al., 2019 [58] for this PRR described in the *Brachypodium* genome, which presented a greater phylogenetic relationship to *At-LYK5*. In clade IV, the Ca1 and 2 *LYP* grouped, in addition to other homologs, to *Mm-LYP1*. The *Mm-LYP1* is a receptor described in white mulberry, besides having a high affinity for chitin, it displays a significant increase in transcriptional profiles in fruits and leaves of mulberry infested with popcorn disease. The *Mm-LYP1* interacts with *Mm-LYK2*, a homolog of *At-CERK1*, present in clade I and grouped with the candidate sequences for *CERK1* in *C. arabica*. The *Mm-LYK2* does not have a high affinity for chitin, but it functions as a co-receptor with intracellular kinase for the PTI signaling [31]. Additionally, in this clade, the *Hv-CEBiP* in barley, has been described for recognizing chitin oligosaccharides derived from *Magnaporthe oryzae* [28] and *Mt-LYM2*, in *Medicago truncatula*, demonstrated specific binding to biotinylated N-acetylchitooctaose in a similar way to *CEBiP* in rice [23,63]. Thus, the receptors cited for the phylogenetic groupings of this study reinforces the possible role of candidate sequences in *C. arabica* as PAMP receptors.

### Transcriptional response of candidate receptors in *C. arabica*

The PAMPS are defined as highly conserved molecules from microorganisms and, therefore, have an essential function in their survival or fitness [73,74]. It is suggested that since PAMPs are essential for the viability or lifestyle of microorganisms, it is less likely that they avoid host immunity through mutation or deletion in these regions [15,75]. Chitin is a PAMP present in the fungal cell wall. Fragments of N-acetylquitooligosaccharides are released by the breakdown of this PAMP by plant chitinases during plant-fungus interactions. These fragments serve as elicitors for the innate immunity of plants by modifying the transcriptional levels of PRRs [23].

In this study, the expression increases were detected from 6 hpi, showing that all candidate PRR were stimulated after the inoculation of *H. vastatrix*. The highest averages of expression were observed at 24 hpi, for most receptors, followed by a decrease at 48 hpi (Fig 5). These results describe an initial stimulus with subsequent suppression. The experiments showed that at 24 hpi it is already possible to detect the penetration of the hypha produced by the appressorium of *H. vastatrix* in stomata of coffee leaves, both in resistant and susceptible genotypes and at 48 hpi the presence of haustoria is already observed [76–78]. In addition, a LRR receptor-like kinase described in this pathosystem has a peak expression at 24 hpi in compatible and incompatible interactions [79], thus suggesting that the signal exchange between the two organisms is already occurring in this period.

To inhibit PTI, some fungal pathogens secrete proteins containing LysM motifs that compete with plant receptors [80,81]. These proteins seem to impede the detection of chitin polymers or interfere with the functioning of essential molecules in the downstream signaling of basal immunity. It is assumed that the decrease in PRR expression in *C. arabica* leaves, observed at 48 hpi, may be related to the suppression of PTI signaling. Fungal effectors such as *Ecp6, ChELP1/2* bind to chitin oligosaccharides released by the action of chitinases and prevent their recognition by the host PRR [80,82], while effectors like *Avr4* protect chitin from fungal cell walls from degradation by host chitinase [83]. In addition, a study of the *H. vastatrix* secretome showed that effector candidates expressed in incompatible interaction (resistance) were more abundant within 24 hours, suggesting that these pre-haustorial effectors could be involved in the attempt to suppress PTI [84].

The expression results of the candidate receptors did not show difference in profiles between the groups of resistant and susceptible cultivars. Despite the IP showing high levels of expression at 24 hpi for the transcripts *Ca1-LYP, Ca2-LYK5* and *Ca-LYK4*, the susceptible cultivar MN showed equivalent levels of expression for *Ca1-CERK1* and *Ca2-LYP* or MN and CV showed comparable levels or even larger than the AR resistant cultivar for *Ca2-CERK1, Ca2-LYP, Ca1-LYK5* and *Ca-LYK4* (Fig 5). This result was expected, since the basal immunity is characterized by being broad-spectrum and non-specific [13,18]. The resistance of coffee to rust has been reported as pre-haustorial [78,85], in which resistant genotypes cease the growth of the fungus with mechanisms of pathogen recognition by resistance proteins. Thus, the difference between resistant and susceptible cultivars is generally evidenced in studies of expression of genes involved in pathogen-specific pathways and not in broad-spectrum receptors, such as PRRs [85].

Additionally, the recognition and signaling of PAMPs occurs when PRRs associate and act as part of multiprotein immune complexes on the cell surface [86,87]. Although they share common structural characteristics, these receptors are distinct in terms of recognized expression patterns and epitopes [24,26,53,63]. This shows that the receptors roles appear to have evolved independently in different groups of plants [26,72]. Therefore, considering that all candidate receptors in coffee, described in this study, increased their expression from 6 hpi in all evaluated cultivars, each one may have possible roles in the basal immunity of *C. arabica*.

## Conclusion

The results indicate that candidate sequences in *C. arabica* have protein domains and motifs characteristic of fungal PRRs and are homologous to *At-CERK1, At-LYK4, At-LYK5* and *Os-CEBiP*. Additionally, the expression of these genes was increased after the inoculation of *H. vastatrix* at all times and cultivars evaluated. Therefore, this study presents an advance in the understanding of the basal immunity of this species.

## Acknowledgments

The authors would like to acknowledge the M.Sc. Antonio Carlos da Mota Porto for his help with the statistical analysis.

## Supporting information

**S1 Fig. Germination of *H. vastatrix* spores observed by optical microscope after 48 hours of inoculum preparation**.

**S2 Fig. Symptoms and signs of *H. vastatrix* in *C. arabica* seedlings**.

(A, B, C, D) Cultivar Mundo Novo IAC 367-4, (E, F) Catuaí Vermelho. (A) abaxial face 20 days after inoculation of the pathogen, (B) adaxial face 20 days after inoculation, (C, E) abaxial face 35 days after inoculation, (D) adaxial face 35 days after inoculation.

**S3 Fig. Stability ranking of the reference genes *14-3-3, GAPDH, EF1a* and *24S* obtained by RefFinder tool**.

(A) Experiments 1, (B) Experiment 2. GM: Geometric mean of the weights from algorithms Delta-Ct, BestKeeper, NormFinder e geNorm.

**S4 Fig. Alignments of CDS from candidate sequences to *CERK1* (*Ca1-CERK1* Scaffold_539**.**592 e Scaffold_1805**.**113)**. The alignments were obtained by CLC Genomics Workbench software. Gray bars show the conservation level of the positions; red letters, the different nucleotides; and red dashes, the gaps. Identity: 71, 33%.

**S5 Fig. Alignments of CDS from candidate sequences to *CERK1* (*Ca2-CERK1* (Scaffold_2193**.**164 e Scaffold_476**.**38) in *C. arabica***. The alignments were obtained by CLC Genomics Workbench software. Gray bars show the conservation level of the positions; red letters, the different nucleotides; and red dashes, the gaps. Identity: 98,28%.

**S6 Fig. Alignments of CDS from candidate sequences to *LYK4* (Scaffold_612**.**376 e Scaffold_952**.**320) in *C. arabica***. The alignments were obtained by CLC Genomics Workbench software. Gray bars show the conservation level of the positions; red letters, the different nucleotides; and red dashes, the gaps. Identity: 98,45%.

**S1 Table. BLASTp analysis of the PRR reference sequences against the *C. arabica* genome in Phytozome**.

